# Obtaining super-resolved images at the mesoscale through Super-Resolution Radial Fluctuations

**DOI:** 10.1101/2024.07.31.606008

**Authors:** Mollie Brown, Shannan Foylan, Liam M. Rooney, Gwyn W. Gould, Gail McConnell

## Abstract

Super-resolution microscopy overcomes the diffraction limit of light to achieve higher spatial resolutions than are typically available when using light microscopy techniques. However, these methods are usually restricted to imaging a very small field of view (FOV). Here, we have applied one of these super-resolution techniques, Super-Resolution Radial Fluctuations (SRRF) in conjunction with the Mesolens, which has the unusual combination of a low-magnification and high numerical aperture, to obtain super-resolved images over a FOV of 4.4 mm x 3.0 mm. We assessed the accuracy of these SRRF images through error maps calculated using a secondary analysis method, Super-resolution Quantitative Image Rating and Reporting of Error Locations (SQUIRREL). We demonstrate it is possible to achieve images with a resolution of 446.3 ± 10.9 nm, providing a ∼1.6-fold improvement in spatial resolution over a uniquely large field, with consistent structural agreement between raw data and SRRF processed images.

**Motivation:** Current super-resolution imaging techniques allow for a greater understanding of cellular structures however they are often complex or only have the ability to image a few cells at once. This small field of view may not represent the behaviour across the entire sample and the manual selection of which restricted ROI to use may introduce bias. Currently, this is often circumvented by stitching and tiling methods which stitch many small ROI together, however this can result in artefacts across an image which poses an issue when analysing data. To combat this, we have used the Mesolens alongside Super-Resolution Radial Fluctuations analysis, to obtain super-resolved images over a field of view of 4.4 mm x 3.0 mm with minimal error.

## Introduction

The spatial resolution achievable in light microscopy discerns the smallest distance at which two individual structures can be resolved. This can be approximated using Rayleigh resolution limit^1^, *r* and is dependent on the numerical aperture, *N*.*A*., of the lens used. (1)

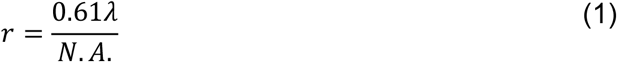

Super-resolution microscopy refers to the group of techniques which surpass this limit to achieve higher resolutions than theoretically possible^2^. Whilst these techniques allow greater understanding of cellular structure through the resolution of finer structures, there are often limitations associated. Methods such as Stimulated Emission Depletion microscopy^3^ and Structured Illumination Microscopy^4,5^ can significantly increase the resolution, but require complex hardware. Single Molecule Localisation Microscopy methods (SMLM) such as Photoactivated Localisation Microscopy^6,7^ and Stochastic Optical Reconstruction Microscopy^8^ are accessible with more conventional microscopes, but these methods require specialised fluorophores, with distinct ON and OFF states. Large datasets capturing these individual states can be reconstructed to produce a super-resolved image, but the fluorophores can be expensive or inaccessible and thousands of images are necessary for reconstruction^9,10^.

Additionally, for both shaped illumination super-resolution and SMLM techniques, a high NA lens is needed and hence the field of view (FOV) is often small, typically less than 100 μm x 100 μm^10,11^ sampling information from only a small number of cells in a single image. As such, manually selected regions of interest (ROIs) may introduce bias and may not be representative of the wider specimen. Stitching and tiling methods have been applied to produce super-resolved images over larger fields^12,13^ but this computational approach can introduce artefacts where the edges of the tiles are poorly matched, or where there is inconsistent illumination or fluorescence across separate tiles. Alternative methods using chip-based illumination have also been reported, achieving resolutions of up to 70 nm, but again the FOV is restricted to 0.5 mm x 0.5 mm^14^.

In recent years, computational methods to produce super-resolved images have been developed which do not require complex equipment or labelling techniques. These methods use a similar principal to SMLM to reconstruct super-resolved images from the intensity fluctuations within diffraction-limited datasets^15–18^. One of the most established methods is SRRF (Super-Resolved Radial Fluctuation)^15^. SRRF utilises both the spatial and temporal information available within standard diffraction-limited widefield microscopy datasets to obtain super-resolved images by following similar principles as SMLM reconstruction without the need for specialised fluorophores. By measuring the radial symmetry (radiality) of the intensity surrounding each pixel in an image, SRRF works to calculate the probability of each pixel containing fluorescence. If a pixel has a uniform radial symmetry (i.e., high radiality) then it is likely fluorescent, while pixels with low radiality may be attributed to noise or spurious non-fluorescent events. The correlation of each point throughout the stack is processed using a selected temporal analysis algorithm, to create a final SRRF image that considers both the spatial and temporal information. There are a variety of adjustable parameters which allow for refinement of any SRRF analysis. Of these, two are particularly impactful on the work shown here and should be noted – the radiality magnification and ring radius. The ring radius determines the radius of the ring which the radiality measurements are taken from and should be adjusted depending on the density of the datasets. When SRRF calculates the radiality for each pixel, it can do this on a singular pixel, or more typically it magnifies each pixel into a grid of *n* × *n* subpixels, where *n* is the magnification, and performs this radiality measurement on each sub-pixel. This often leads to resolution up to five-fold higher in the final reconstruction^16^, however increasing the pixels within the image also in turn increases the size of the image file.

Following SRRF processing it is important to verify the accuracy of the super-resolution reconstructions. This is achieved using two primary methods. The first is analysis with super-resolution quantitative image rating and reporting of error locations (SQUIRREL)^19^, to both map the accuracy across the image and quantify the error within the super-resolution images using Resolution Scaled Pearson’s coefficient (RSP) and Resolution Scaled Error (RSE). The second method is through measuring the resolution of the final super-resolution image with image decorrelation analysis^20^.

SRRF has been widely used for a variety of biological applications, with studies reporting an increase in spatial resolution from 235 nm to less than 100 nm in the cell wall of xylem from Douglas Fir trees, although over a limited field with only a few cells analysed per image^21^. This approximate 2-fold improvement in resolution is consistent across other work using SRRF, including the quantification and localisation of azurophilic granules in neutrophil leukocytes^22^, where the resolution of raw data was increased from 160 nm to better than 100 nm using SRRF but, again, only a few cells were captured in each image^22^.

Here, we demonstrate that by applying SRRF, in conjunction with diffraction-limited widefield Mesolens imaging, it is possible to obtain super-resolved images over a large multi-millimetre FOV through the analysis of intensity fluctuations across widefield Mesolens images.

The unusual combination of low magnification and high NA from the Mesolens (4x, 0.47 NA) enables imaging over a FOV of 4.4 mm x 3.0 mm, with a lateral resolution of up to 700nm^23^ and the ability to capture over 700 cells per image^24^. Using SRRF, we aimed to improve the lateral resolution as far as possible while retaining the large FOV for super-resolved imaging of larger cell populations than are traditionally imaged using SRRF microscopy.

## Results

Figure 1A shows a full Mesolens FOV image of tubulin-labelled HeLa cells, following SRRF processing, demonstrating the scale and the volume of cells captured per image, along with the absence of any stitching and tiling artefacts across the entire FOV of super-resolved reconstruction. Digital zooms of ROI are highlighted to demonstrate the improvement in contrast and resolution between the original Mesolens images (Figure 1, B-D) and the corresponding SRRF equivalent (Figure 1, E-F). Between each of the corresponding ROI pairs there is a clear improvement in image clarity and spatial resolution, allowing for a level of detail not previously visualised across the mesoscale. No additional image processing was carried out on either set of images.

**Figure 1:**
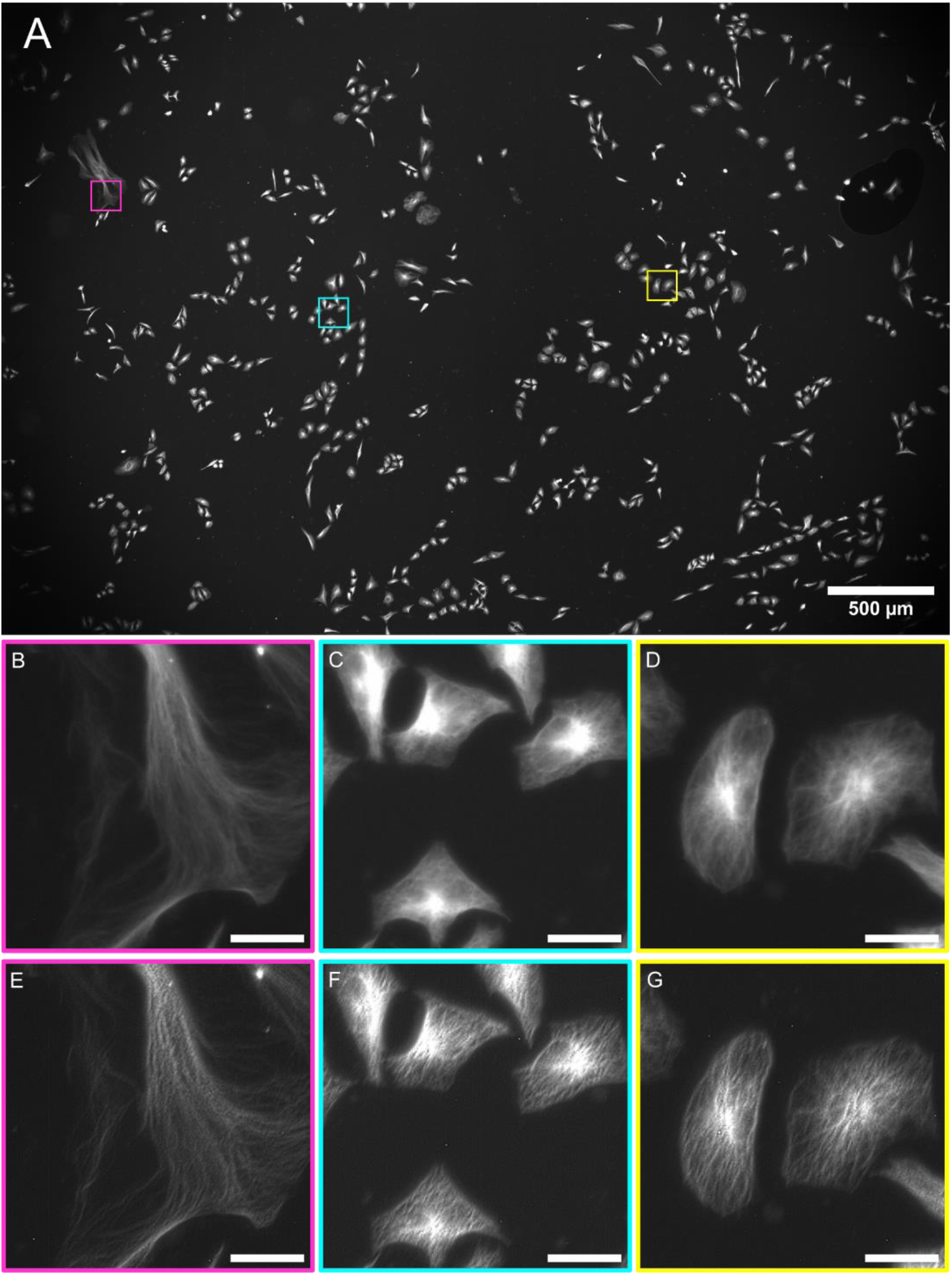
SRRF processing of Mesolens epifluorescence diffraction limited images of tubulin in HeLa cells. (A) The complete Mesolens FOV showing the SRRF processed image of the tubulin in HeLa cells labelled with AF488 with three highlighted ROI; magenta, cyan, and yellow. (B-D) A digital zoom of the raw widefield epifluorescence diffraction limited Mesolens image is shown for each ROI. Scale bars are 30*μm* for all ROIs. (E-G) A digital zoom of the improved ROI following SRRF processing. Scale bars are 30*μm* for all ROIs.

SQUIRREL analysis on the data set shown in Figure 1 is presented in Figure 2. These data show there is minimal error and high levels of agreement between the original and the super-resolution images across the full FOV (Figure 2A). The same ROIs in Figure 1 are shown with SQUIRREL processing in Figure 2, B-D. These show the areas of disagreement between the raw Mesolens image, and the super-resolution equivalent. These ROIs highlight that this low error and high agreement is continuous across both the full FOV and the individual regions, and that while there are areas of disagreement across the map, the disagreement is small. Figure 2, E-G show digital zooms of a contrast adjusted error map. The image was adjusted using Contrast Limited Adaptive Histogram Equilization (CLAHE) with the default parameters (blocksize = 127, histogram bins = 256, maximum slope= 3.0, mask = none, fast = false)^25^, to highlight the areas of discrepancy. We note these discrepancies are largely confined to areas where the image is saturated.

**Figure 2:**
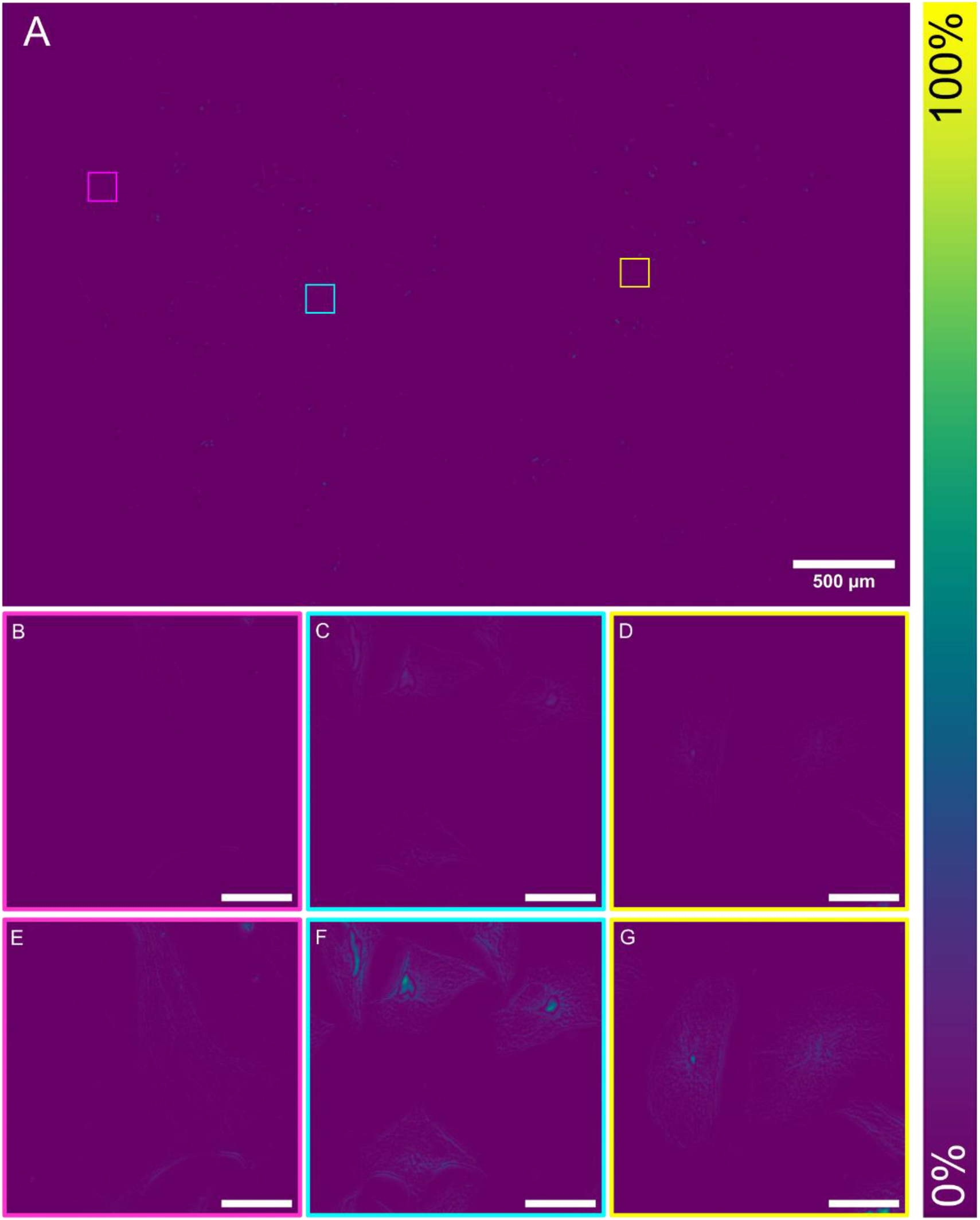
SQUIRREL processing of SRRF processed Mesolens data. (A) The reconstructed SQUIRREL error map showing areas of agreement between the raw widefield epifluorescence diffraction limited Mesolens image and the SRRF reconstruction. (B-D) In the middle row a digital zoom of the error map is displayed for each ROI. Scale bars are 30 *μm* for all ROIs. (E-G) A contrast adjusted ROI below (E-G), highlighting the areas of error within each ROI. Scale bars are 30 *μm* for all ROIs.

For the dataset shown in Figures 1 and 2, this high fidelity between images is confirmed by the RSP value of 0.989 ± 0.004. RSP is a normalised value between -1 and 1, with 1 representing structurally identical images. The value shown here indicates high structural agreement, and minimal artefacts between both images, alongside the intensity dependent RSE result of 1.13 % ± 0.23 %. The RSE value presented has been normalised against the total intensity range in the SQUIRREL error map, allowing for simplified comparisons between different datasets. The resolution of this super-resolution reconstruction in Figure 1A was also measured from three ROI at 442.9 nm (See Figure S1A-1C). A significant improvement in image resolution, which surpasses the theoretical maximum resolution achievable on the Mesolens – providing super-resolved images.

To calculate an average resolution measurement achieving using this method, measurements were repeated for three biological repeats of HeLa cells where tubulin was labelled with an AF488 antibody conjugate. There was an average RSP value of 0.991 ± 0.006, and an average RSE value of 0.81 % ± 0.30 %, with consistently low error rates and high agreement across different images. An average resolution was also recorded using three ROI from each sample and measured at 446.3 ± 10.9 nm (See Figure S1).

Due to the size of the Mesolens data it is currently not feasible to increase the SRRF magnification above 2 on raw data without surpassing the maximum allowed file size for processing. For higher magnification values SRRF processing is possible on ROIs only, so a magnification of 2 is used here to allow for processing of raw Mesolens data without the addition of stitching and tiling artefacts. To demonstrate that using a higher magnification would have minimal impact on image resolution, SRRF was carried out on the same ROIs as shown in Figure 1, using increasing magnification and the average resolution of these ROI for each magnification was calculated, as shown in Figure 3.

**Figure 3:**
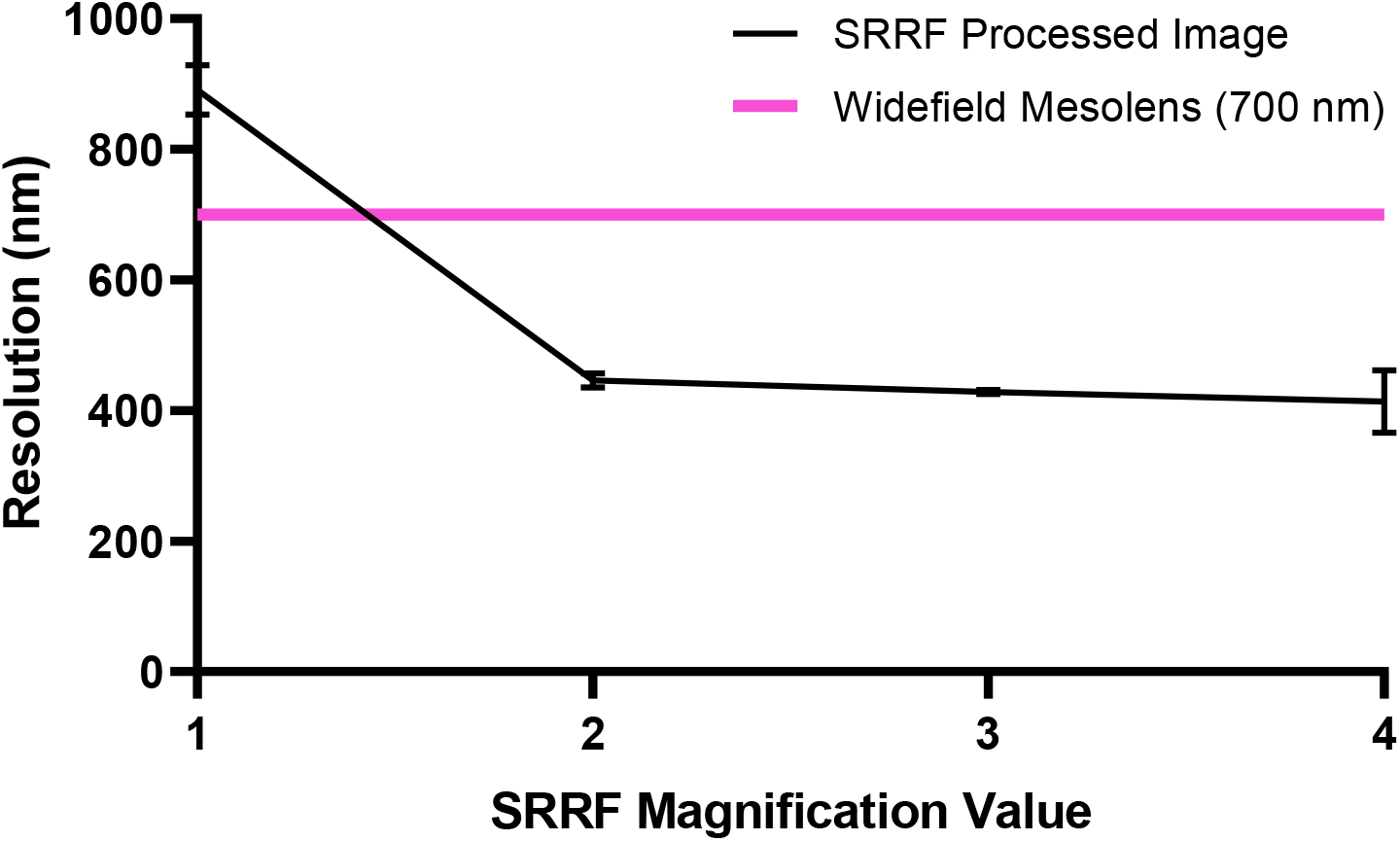
The image resolution measured using image decorrelation analysis, for increasing SRRF magnification, comparative to the previously shown Mesolens resolution of 700 nm^23^. Error bars represent standard deviation.

### Applications to non-filamentous structures

To demonstrate this method is not confined to filamentous structures, Figure 4 shows a full Mesolens FOV capture of GFP-tagged GLUT4 glucose transporters in 3T3-L1 fibroblasts following SRRF processing, with digitally magnified ROIs to again show the improvement in the SRRF reconstruction from the original widefield epifluorescence diffraction limited Mesolens image. Figure 4A shows the entire FOV of super-resolved reconstruction, and the intact cells captured again without any stitching and tiling artefacts. Digital zooms of ROIs demonstrate the improvement in contrast and resolution between the original Mesolens images (Figure 4, B-D) and SRRF equivalent (Figure 4, E-F). Again, between each of the ROI pairs there is a clear improvement in resolution and image clarity, allowing for clearer visualisation of the GLUT4 molecules throughout the cell interior. GLUT4 is sequestered within multiple different intracellular compartments in the absence of insulin^26,27^, and the images shown here provide a clear illustration of this distribution. The resolution of this image was measured from three ROI at 431.6 ± 9.8 nm (See Figure S2), which again surpasses the maximum optical resolution theoretically possible on the Mesolens, obtaining super-resolved images.

**Figure 4:**
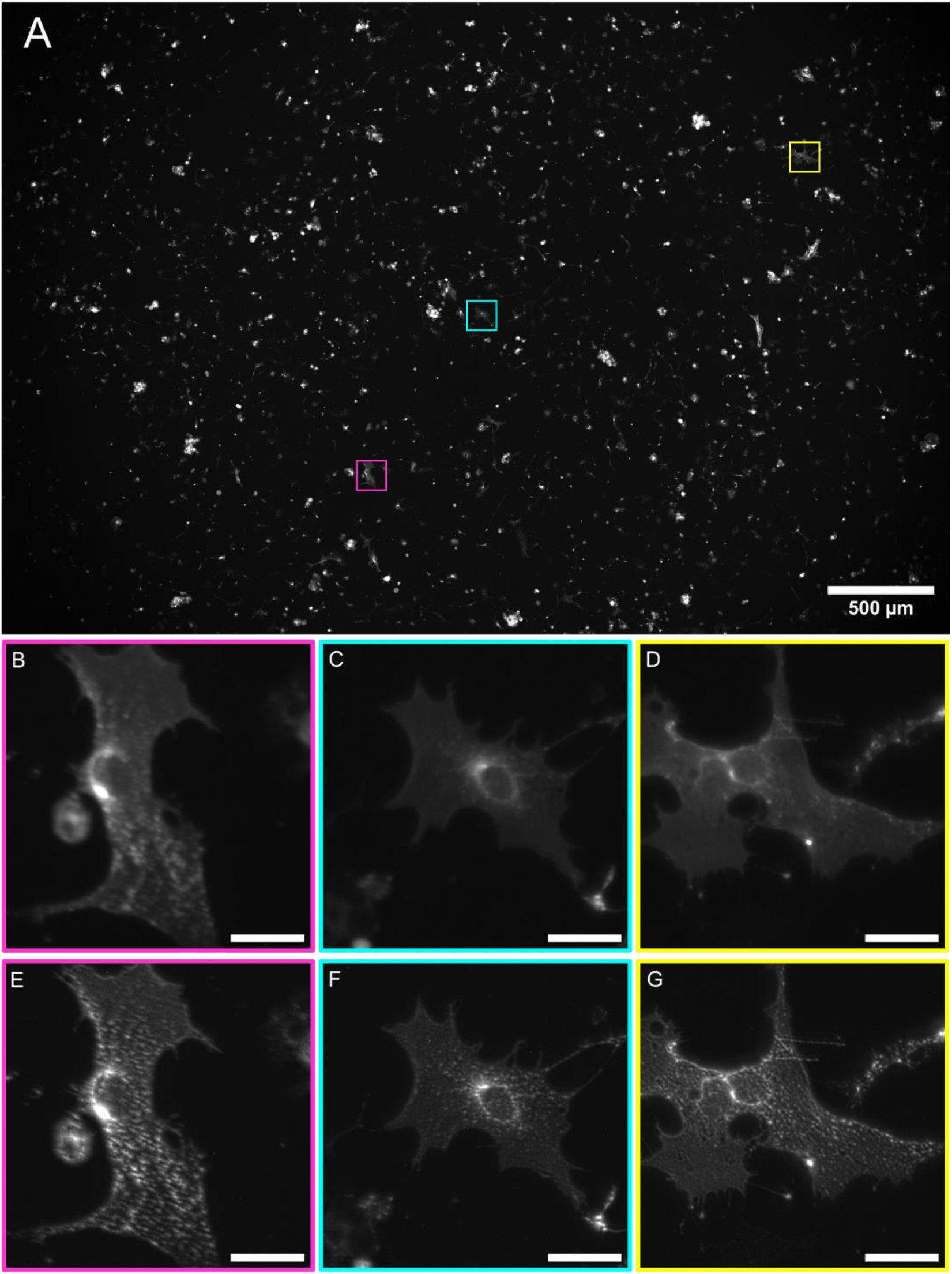
SRRF processing of Mesolens epifluorescence diffraction limited images of GFP-expressing GLUT-4 in 3T3-L1 adipocytes. (A)The complete Mesolens FOV showing the SRRF processed image of the GLUT-4 in 3T3-L1 adipocytes expressing GFP, with three highlighted ROI; magenta, cyan, and yellow. (B-D) A digital zoom of the raw widefield epifluorescence diffraction limited Mesolens image is shown for each ROI. Scale bars are 30 *μm* for all ROIs. (E-G) A digital zoom of the improved ROI following SRRF processing. Scale bars are 30 *μm* for all ROIs.

SQUIRREL analysis on this dataset is shown in in Figure 5. There is again minimal error and high levels of agreement between the raw widefield epifluorescence diffraction limited Mesolens image and the SRRF reconstruction throughout across the full FOV (Figure 5A). Digital zooms of the ROIs (Figure 5, B-D) highlight that there is some error present across the FOV, but it is minimal. Figure 5, E-G show digital zooms of a contrast adjusted error map. The image was adjusted using CLAHE with the default parameters (blocksize = 127, histogram bins = 256, maximum slope= 3.0, mask = none, fast = false)^25^, to better highlight the areas of discrepancy. For this dataset the average RSP value was calculated as 0.993 ± 0.003, and the RSE value as 0.68 % ± 0.14 %, again demonstrating the high fidelity and low levels of error between the original raw diffraction limited widefield Mesolens image, and the SRRF processed reconstruction.

**Figure 5:**
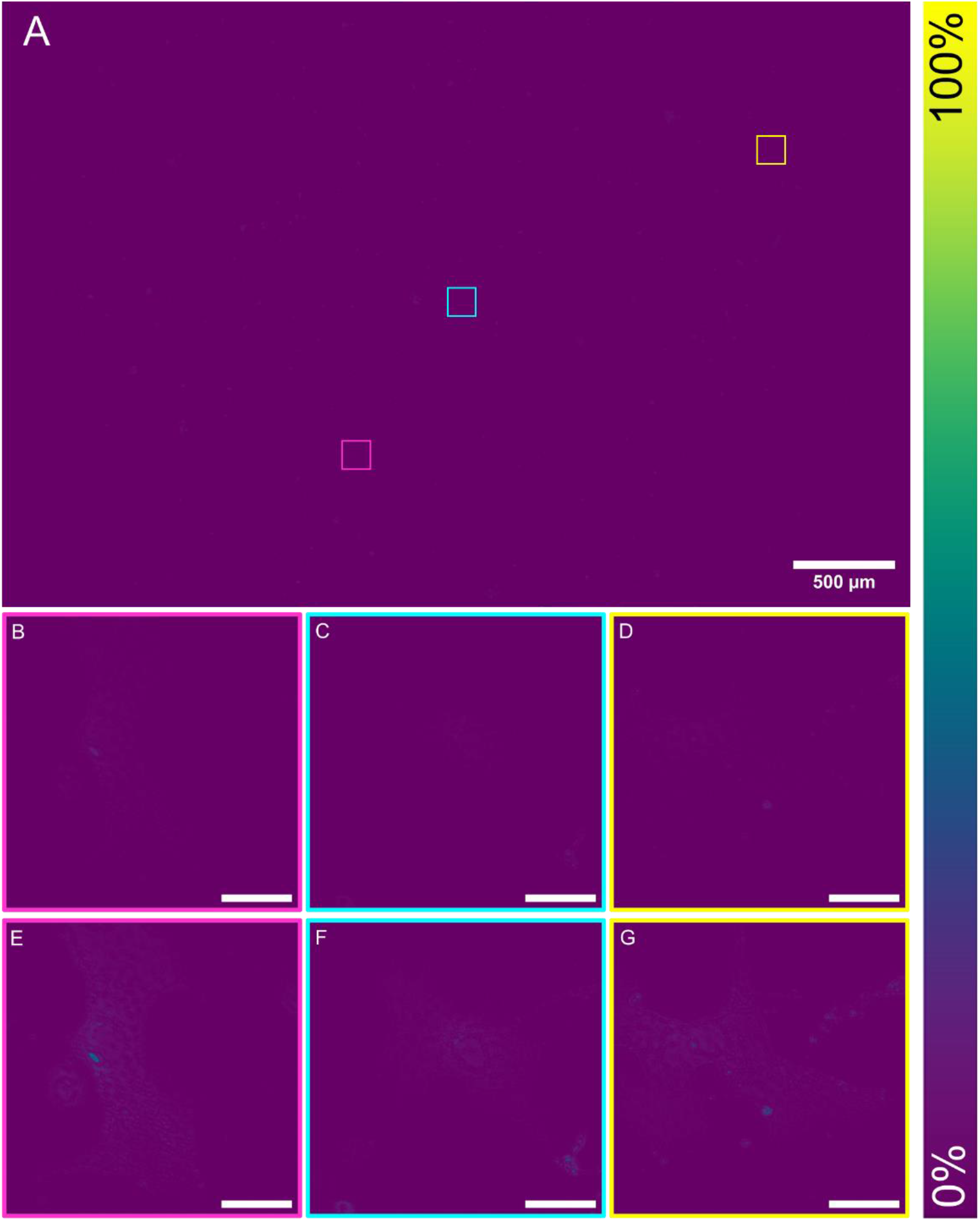
The reconstructed SQUIRREL error map showing areas of agreement between the raw widefield epifluorescence diffraction limited Mesolens image and the SRRF reconstruction. (A) The error map for the complete FOV of the SRRF processed Mesolens image with highlighted ROI; magenta, cyan and yellow. (B-D) A digital zoom of the error map is displayed for each ROI. Scale bars are 30*μm*. (E-G) A contrast adjusted ROI highlighting the areas of error within each ROI. Scale bars are 30*μm* for each ROI.

**Figure 6:**
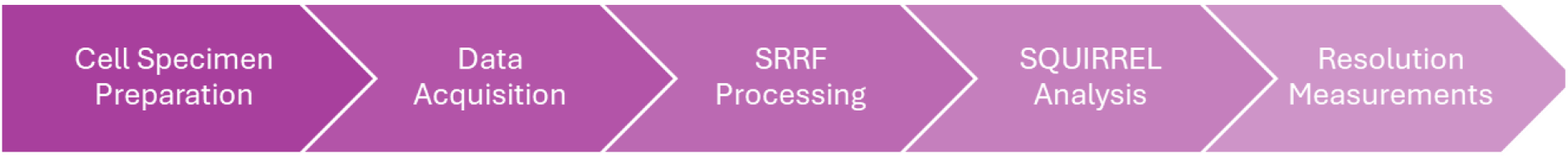
An overview of the processing stages in obtaining super-resolved Mesolens images.

## Discussion

Following SRRF processing there is a clear visual improvement in spatial resolution and contrast, allowing for clearer visualisation of the filamentous and non-filamentous structures imaged. Using the 446.3 nm resolution obtained from the SRRF (M=2) value for filamentous structures and a wavelength of 530nm with equation (1), we can consider that the effective NA of the Mesolens has increased from 0.47 to approximately 0.72, with no reduction in FOV.

The error maps show that there is minimal error and that the SRRF images are in high agreement with the original Mesolens images. It is possible to see that these are largely within the high-intensity areas where it is difficult to visibly discern fine structures within the original images. This low error is also validated by the average RSP and RSE values, which indicates there is high fidelity between both images. This is further illustrated by the resolution measurement, which show a notable improvement following SRRF, surpassing the resolution limit for the Mesolens, and achieving super-resolved images.

As seen in Figure 4, this method is not limited to filamentous structures and can be used to accurately improve the resolution of other, non-filamentous cellular structures, demonstrating its capabilities. The resolution of the 3T3-L1 fibroblasts expressing GFP labelled GLUT4 was measured at 431.6 ± 9.8 nm, which is higher than the achievable resolution stated previously. This resolution of 446.3 ± 10.9 nm is calculated from three ROIs from three biological replicates (compared to the three ROIs from one biological replicate here) and as such provides a theoretical achievable resolution, which may range depending on the samples and the imaging conditions used. The intracellular localisation of GFP-tagged GLUT4 to vesicles within the cell is clearly revealed using this approach.

Using SRRF in conjunction with the Mesolens allows for an increased understanding of many cellular structures, with an improvement in resolution comparable to other work utilising SRRF. However, using this method circumvents any potential artefacts introduced by stitching and tiling, or from manually selecting ROIs – providing a more accurate understanding of the entire field. It has a wide range of potential applications - the increase in spatial resolution would allow for more accurate co-localisation studies, to better determine the location and interaction of molecules throughout a cell population. The use of widefield epifluorescence imaging in obtaining these images is also less harsh to samples than other conventional super-resolution microscopy methods, allowing for preservation for repeated imaging.

Other work utilising SRRF often uses higher magnification values to obtain super-resolved images^21,28,29^. Increasing the magnification also increases the size of the image being processed, as each original pixel is split into a grid of smaller pixels, increasing the data contained within one image. This increase in image size limits the processing of intact Mesolens data to using a magnification of 2 as beyond this processing is only possible on ROIs. As seen in Figure 3 there is a slight increase in the resolution achievable when increasing magnification beyond 2. However, carrying out processing on individual ROIs would introduce stitching and tiling artefacts, and the impact of this would outweigh any marginal improvements in resolution. If future work was able to circumvent the computational limitations currently in place it may allow for the use of higher magnifications to provide an additional increase in resolution. The computational power available here also limits the processing time, particularly for error analysis using SQUIRREL, if this can be increased then processing time would be significantly reduced.

A recent SRRF successor, Enhanced Super-Resolution Radial Fluctuations (eSRRF)^17^, expands upon the same analysis method to improve the accuracy of the super-resolution images, and further increase the achievable resolutions. However, due to the size of the Mesolens datasets used, it was deemed impractical to use eSRRF. It would be possible to split the Mesolens images into individual tiles and apply eSRRF to each tile, however, due to the radiality component of the method, the pixels at the edge of each tile could not be properly analysed, with only a fraction of the radial information available. This would introduce artefacts and inconsistent intensities across the tile edges. Due to this, SRRF is used rather than eSRRF for this study.

Although both SRRF and its successor eSRRF are capable of live cell super-resolution reconstructions, due to the long acquisition time of one Mesolens image, 13.5 s when using a 1500 ms exposure time, and the current absence of an environmental imaging chamber this method is unsuitable for the imaging of fast cell processes. However, there may be scope for future super-resolution live-cell work on the Mesolens with new sensors with higher pixel numbers and faster chip-shifting.

In this work we have demonstrated using SRRF in conjunction with diffraction-limited widefield Mesolens images, that it is possible to obtain super-resolved images at the mesoscale – achieving a resolution of 446.3nm across a 4.4 mm X 3.0 mm FOV. With SQUIRREL analysis and error maps demonstrating consistent agreement between the original and SRRF processed images validating the accuracy of the reconstructions. This provides a comparatively cost-efficient and simple method for obtaining accurate super-resolution images over a large FOV, allowing for a simultaneous understanding of the subcellular structures alongside their large-scale interactions.

## Acknowledgements

M.B. is supported by a Research Excellence Award Studentship from the Engineering & Physical Sciences Research Council. S.F., G.W.G and G.M. were funded by the Biotechnology and Biological Sciences Research Council, BB/X005178/1. L.M.R. was funded by the Leverhulme Trust. G.M. was funded by The Medical Research Council, MR/K015583/1, and the Biotechnology and Biological Sciences Research Council, BB/P02565X/1 and BB/T011602/1.

## Author Contributions

Methodology, M.B.; Investigation, M.B., S.F. and L.R.; Resources, M.B., S.F., L.R.; Writing – Original Draft, M.B.; Writing – Review & Editing, S.F., L.R., G.G., and G.M.; Conceptualisation, M.B., L.R., and G.M.; Supervision, G.G. and G.M.; Funding Acquisition, G.G. and G.M.

## STAR Methods

### Key Resources table

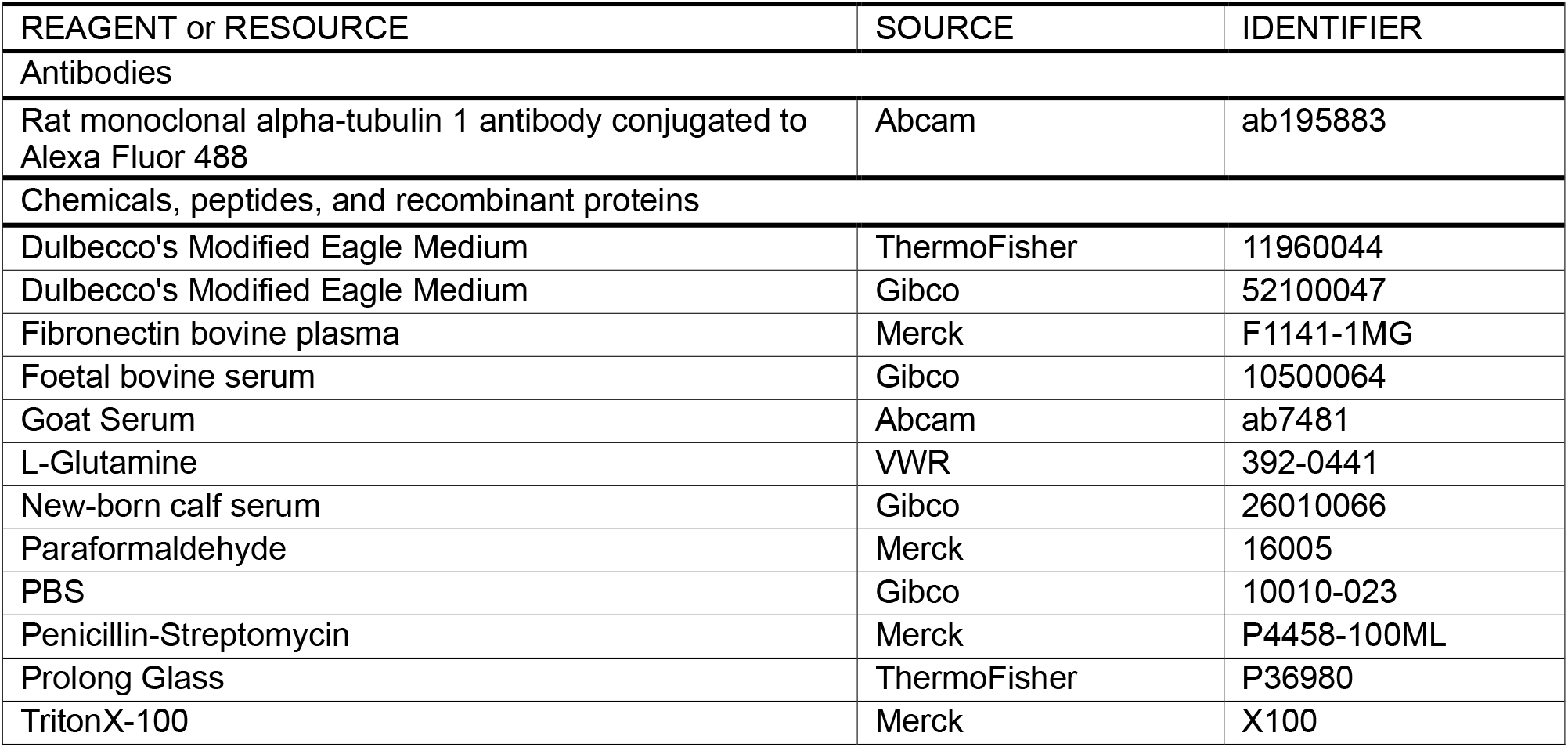

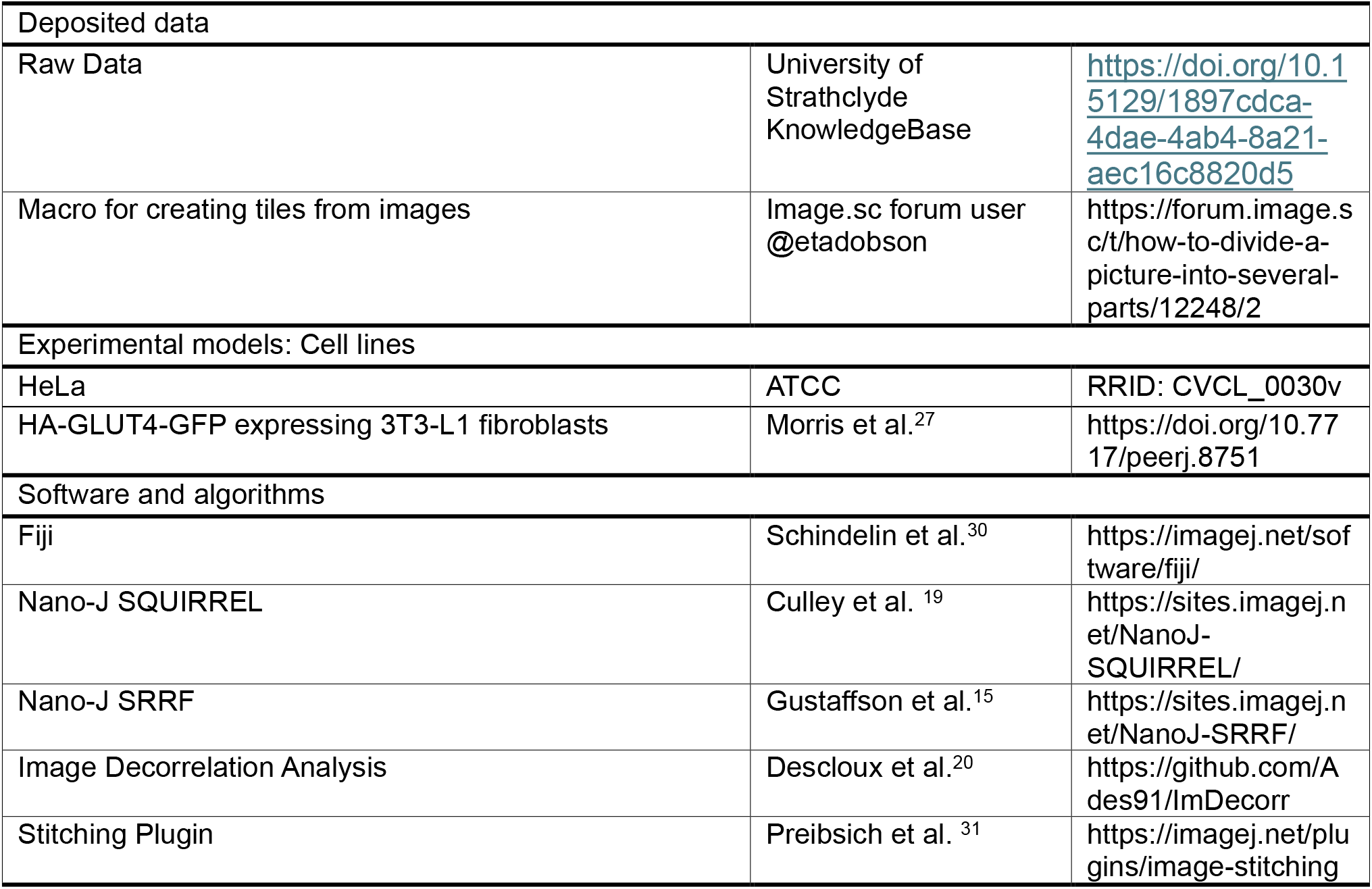

### Resource availability

#### Lead Contact

Further information and requests for resources and should be directed to and will be fulfilled by the lead contact, Mollie Brown.

## Materials availability

This study did not generate new unique reagents.

## Data and code availability

All data underpinning this publication are publicly available on request from the University of Strathclyde KnowledgeBase at: https://doi.org/10.15129/1897cdca-4dae-4ab4-8a21-aec16c8820d5. This dataset includes raw images in .tiff format for all imaging with accompanying ROI sets.

### Experimental model and study participant details

HeLa cells were cultured at 37°C, 5% CO_2_, in Dulbecco’s Modified Eagle Medium (DMEM) (ThermoFisher) containing 1% Penicillin-Streptomycin, 1% L-Glutamine, and 10% foetal bovine serum. Coverslips were coated with a 1:500 dilution of fibronectin bovine plasma and incubated for 24 hours prior to fixation.

A stable cell line of HA-GLUT4-GFP expressing 3T3-L1 cells was previously generated in the lab^27^ and plated on 18 mm diameter coverslips and grown in DMEM (Gibco) supplemented with 10% new-born calf serum (NBCS), 5% L-glutamine and 5% penicillin-streptomycin and maintained at 37°C, 10% CO2^32^.

## Method Details

### Cell Specimen preparation

Coverslips containing HeLa cells were fixed with 4% paraformaldehyde (PFA) at 37°C for 15 minutes, before being blocked and permeabilised for 30 minutes at 37°C in an immunofluorescence (IF) buffer comprised of 2.5% Goat Serum and 0.3% TritonX-100 in PBS, labelled using anti-Tubulin rat monoclonal conjugated with Alexa Fluor 488 at a dilution of 1:100 in IF buffer and incubated at 4°C for 24 hours in a dark culture dish with stable humidity. The samples were then mounted using Prolong Glass to reduce photobleaching during prolonged imaging and left to set for at least 24 hours prior to imaging.

### Non-Filamentous Structures

Following differentiation into adipocytes cells were put on ice, washed once with ice-cold 1X PBS and fixed in 4% PFA for 30 minutes at room temperature. Coverslips were washed three times with 1X PBS before mounting on a microscope slide with Prolong Glass. Specimens were allowed to set fully prior to imaging.

### Mesolens Imaging and Data Acquisition

Diffraction-limited images were obtained with the Mesolens using the widefield fluorescence illumination modality, a diagram of this system set-up is shown in Figure 7. Specimens were illuminated using a 490 ± 20 nm LED sourced from a multi-wavelength illuminator (pE-4000, CoolLED), and fluorescence emission was detected at 525 ± 25 nm. The maximum power of the excitation source at the specimen plane was 35.7 ± 0.1 mW. Imaging was carried out with the Mesolens correction collars set for water immersion to minimise spherical aberration. To ensure statistically independent images for SRRF processing, a time-lapse series of the fixed samples was taken to obtain 40 images, with the interval between image capture is typically set to 2 – 3 seconds as the samples are fixed. Other work utilising SRRF can use up to hundreds of individual images^29^ to produce the final reconstruction, however, due to the size of the Mesolens image datasets and the computational restrictions this entails the datasets here are limited to 40 images for SRRF processing. This restricted dataset may reduce the ability of SRRF processing to supress noise, to avoid this exposure time was typically kept above 1500 ms when imaging, as longer exposure times would result in a higher SNR.

**Figure 7:**
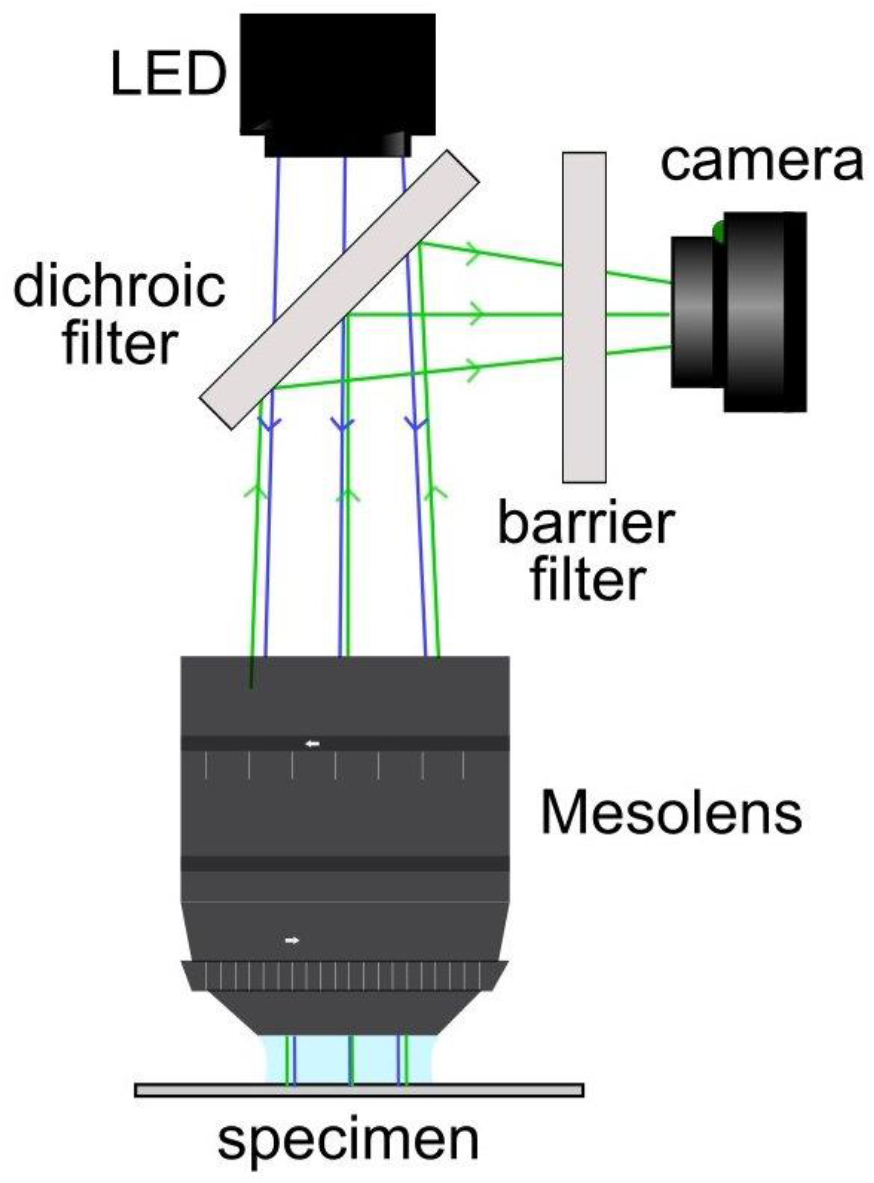
A schematic diagram of the widefield epifluorescence Mesolens system.

To ensure the diffraction-limited widefield Mesolens images captured are of the highest resolution, images are captured using a camera with a chip-shifting sensor (VNP-29MC, Vieworks). This shifting moves the 29-megapixel chip in a 3-by-3 array, giving an image resolution of 19728 × 13152 pixels for a total 259.5-megapixel capture. Over the 4.4 × 3.0 mm FOV this results in a 226 nm pixel size - satisfying Nyquist sampling^33^. The acquisition of one full FOV image using the sensor shifting camera took 13.5 s when using a 1500 ms exposure. When including the additional time for the transfer of image data to the computer, and the interval between captures, imaging of a 40-image stack typically took approximately 22 minutes.

### SRRF Processing

The diffraction-limited widefield Mesolens images were opened within Fiji^30^. To avoid the possibility of introducing any artefacts no preliminary processing was carried out, instead using the Nano-J SRRF plugin^16^ the raw images can be processed. There were restrictions on the parameters which can be used when processing Mesolens images due to the unconventional file size of the datasets, and as such the parameters used vary slightly from the default values. The ring radius was set to 1.90 - when carrying out quantification of the image resolution this was found to result in the highest resolution whilst also allowing for ease of processing. The radiality magnification was reduced so that the large datasets could be processed. This was kept at 2 as it allowed for SRRF processing of intact Mesolens datasets, if the magnification was increased further processing could only be carried out on small ROIs. To ensure this reduced magnification did not limit the resolution achievable with Mesolens datasets the magnification was increased on smaller ROIs and yielded no significant increase in resolution, shown in Figure 5, keeping the magnification at 2 allowed for high resolutions, but without introducing the artefacts stitching and tiling many small ROIs processed with a higher magnification would. For the temporal analysis, Temporal Radiality Pairwise Product Mean (TRPPM) was selected as the fluorophores used do not blink, but instead have slight intensity fluctuations which TRPPM is best suited to process. Using TRPPM over alternative options such as Temporal Radiality Average is also beneficial as it provides additional noise suppression, which is beneficial given the limitations on the size of the datasets used^15^. Any unmentioned parameters remained at the default value. Typical processing time of one dataset was ∼95 minutes, including opening the dataset within Fiji. The computational power available was limited, and this figure could be greatly reduced in further iterations.

### Quantification and statistical analysis

#### SQUIRREL Analysis and Resolution Measurements

Following SRRF processing the accuracy of the output image was first assessed using SQUIRREL. Due to the size of the processed SRRF images SQUIRREL analysis could not be conducted on the intact image. As there is no radiality component within SQUIRREL, the image was split into tiles and later recombined following processing without the introduction of artefacts on the edge of each tile. Macros were used within Fiji to split both a reference Mesolens diffraction-limited widefield image and super-resolution SRRF image into equal tiles, from here it was possible to perform SQUIRREL analysis on each of the matching tile pairs, with the individual SQUIRREL error tiles then stitched^31^ into one complete error map. The individual RSP and RSE values calculated for each tile were averaged to determine the value for the complete image, with the error taken as the standard deviation. RSP measures the Pearson correlation coefficient between the images on a normalised scale between -1 and 1; with 1 being identical images. This acts as a measurement of the structural correlation between images and is independent of intensity. RSE is an intensity dependant measurement of the root-mean-square error between images, where lower values indicant better agreement between the raw image and the super-resolution equivalent. For ease of comparison between data sets, the RSE value was normalised against the total range of possible values. Typically, each Mesolens image is split into 36 tiles for processing, and SQUIRREL analysis of each pair of tiles takes approximately 1 hour.

The resolution was measured using image decorrelation analysis^20^ Fiji plugin, with the default parameters. Due to restrictions imposed by the large Mesolens file size, the measurements were carried out on n=3 ROIs from each super-resolution reconstruction, and the final resolution for each image is an average of these ROI, there was no additional image-processing prior to this stage to ensure an accurate measurement. The achievable resolution presented here is an average of N = 9 ROI, across N = 3 tubulin labelled datasets – again the error is taken as the standard deviation.

## References

1. Rayleigh (1879). XXXI. Investigations in optics, with special reference to the spectroscope. Lond. Edinb. Dublin Philos. Mag. J. Sci. 8, 261–274. 10.1080/14786447908639684.

2. Schermelleh, L., Ferrand, A., Huser, T., Eggeling, C., Sauer, M., Biehlmaier, O., and Drummen, G.P.C. (2019). Super-resolution microscopy demystified. Nat. Cell Biol. 21, 72–84. 10.1038/s41556-018-0251-8.

3. Hell, S.W., and Wichmann, J. (1994). Breaking the diffraction resolution limit by stimulated emission: stimulated-emission-depletion fluorescence microscopy. Opt. Lett. 19, 780. 10.1364/OL.19.000780.

4. Heintzmann, R., and Cremer, C.G. (1999). Laterally modulated excitation microscopy: improvement of resolution by using a diffraction grating. In, I. J. Bigio, H. Schneckenburger, J. Slavik, K. Svanberg, and P. M. Viallet, eds., pp. 185–196. 10.1117/12.336833.

5. Gustafsson, M.G.L. (2000). Surpassing the lateral resolution limit by a factor of two using structured illumination microscopy: SHORT COMMUNICATION. J. Microsc. 198, 82–87. 10.1046/j.1365-2818.2000.00710.x.

6. Betzig, E., Patterson, G.H., Sougrat, R., Lindwasser, O.W., Olenych, S., Bonifacino, J.S., Davidson, M.W., Lippincott-Schwartz, J., and Hess, H.F. (2006). Imaging Intracellular Fluorescent Proteins at Nanometer Resolution. Science 313, 1642– 1645. 10.1126/science.1127344.

7. Hess, S.T., Girirajan, T.P.K., and Mason, M.D. (2006). Ultra-High Resolution Imaging by Fluorescence Photoactivation Localization Microscopy. Biophys. J. 91, 4258– 4272. 10.1529/biophysj.106.091116.

8. Rust, M.J., Bates, M., and Zhuang, X. (2006). Sub-diffraction-limit imaging by stochastic optical reconstruction microscopy (STORM). Nat. Methods 3, 793–796. 10.1038/nmeth929.

9. van de Linde, S., Löschberger, A., Klein, T., Heidbreder, M., Wolter, S., Heilemann, M., and Sauer, M. (2011). Direct stochastic optical reconstruction microscopy with standard fluorescent probes. Nat. Protoc. 6, 991–1009. 10.1038/nprot.2011.336.

10. Lelek, M., Gyparaki, M.T., Beliu, G., Schueder, F., Griffié, J., Manley, S., Jungmann, R., Sauer, M., Lakadamyali, M., and Zimmer, C. (2021). Single-molecule localization microscopy. Nat. Rev. Methods Primer 1, 39. 10.1038/s43586-021-00038-x.

11. Görlitz, F., Guldbrand, S., Runcorn, T.H., Murray, R.T., Jaso-Tamame, A.L., Sinclair, H.G., Martinez-Perez, E., Taylor, J.R., Neil, M.A.A., Dunsby, C., et al. (2018). easySLM-STED: Stimulated emission depletion microscopy with aberration correction, extended field of view and multiple beam scanning. J. Biophotonics 11, e201800087. 10.1002/jbio.201800087.

12. Ortkrass, H., Schürstedt, J., Wiebusch, G., Szafranska, K., McCourt, P., and Huser, T. (2023). High-speed TIRF and 2D super-resolution structured illumination microscopy with a large field of view based on fiber optic components. Opt. Express 31, 29156. 10.1364/OE.495353.

13. Alvelid, J., and Testa, I. (2020). Stable stimulated emission depletion imaging of extended sample regions. J. Phys. Appl. Phys. 53, 024001. 10.1088/1361-6463/ab4c13.

14. Helle, Ø.I., Coucheron, D.A., Tinguely, J.-C., Øie, C.I., and Ahluwalia, B.S. (2019). Nanoscopy on-a-chip: super-resolution imaging on the millimeter scale. Opt. Express 27, 6700. 10.1364/OE.27.006700.

15. Gustafsson, N., Culley, S., Ashdown, G., Owen, D.M., Pereira, P.M., and Henriques, R. (2016). Fast live-cell conventional fluorophore nanoscopy with ImageJ through super-resolution radial fluctuations. Nat. Commun. 7, 12471. 10.1038/ncomms12471.

16. Culley, S., Tosheva, K.L., Matos Pereira, P., and Henriques, R. (2018). SRRF: Universal live-cell super-resolution microscopy. Int. J. Biochem. Cell Biol. 101, 74– 79. 10.1016/j.biocel.2018.05.014.

17. Laine, R.F., Heil, H.S., Coelho, S., Nixon-Abell, J., Jimenez, A., Wiesner, T., Martínez, D., Galgani, T., Régnier, L., Stubb, A., et al. (2023). High-fidelity 3D live-cell nanoscopy through data-driven enhanced super-resolution radial fluctuation. Nat. Methods 20, 1949–1956. 10.1038/s41592-023-02057-w.

18. Torres-García, E., Pinto-Cámara, R., Linares, A., Martínez, D., Abonza, V., Brito-Alarcón, E., Calcines-Cruz, C., Valdés-Galindo, G., Torres, D., Jabloñski, M., et al. (2022). Extending resolution within a single imaging frame. Nat. Commun. 13, 7452. 10.1038/s41467-022-34693-9.

19. Culley, S., Albrecht, D., Jacobs, C., Pereira, P.M., Leterrier, C., Mercer, J., and Henriques, R. (2018). Quantitative mapping and minimization of super-resolution optical imaging artifacts. Nat. Methods 15, 263–266. 10.1038/nmeth.4605.

20. Descloux, A., Grußmayer, K.S., and Radenovic, A. (2019). Parameter-free image resolution estimation based on decorrelation analysis. Nat. Methods 16, 918–924. 10.1038/s41592-019-0515-7.

21. Donaldson, L.A. (2022). Super-resolution imaging of Douglas fir xylem cell wall nanostructure using SRRF microscopy. Plant Methods 18, 27. 10.1186/s13007-022-00865-3.

22. Johnson, J.L., Meneses-Salas, E., Ramadass, M., Monfregola, J., Rahman, F., Carvalho Gontijo, R., Kiosses, W.B., Pestonjamasp, K., Allen, D., Zhang, J., et al. (2022). Differential dysregulation of granule subsets in WASH-deficient neutrophil leukocytes resulting in inflammation. Nat. Commun. 13, 5529. 10.1038/s41467-022-33230-y.

23. McConnell, G., Trägårdh, J., Amor, R., Dempster, J., Reid, E., and Amos, W.B. (2016). A novel optical microscope for imaging large embryos and tissue volumes with sub-cellular resolution throughout. eLife 5, e18659. 10.7554/eLife.18659.

24. Foylan, S., Amos, W.B., Dempster, J., Kölln, L., Hansen, C.G., Shaw, M., and McConnell, G. (2023). MesoTIRF: A prism-based Total Internal Reflection Fluorescence illuminator for high resolution, high contrast imaging of large cell populations. Appl. Phys. Lett. 122, 113701. 10.1063/5.0133032.

25. Zuiderveld, K. (1994). Contrast limited adaptive histogram equalization. In Graphics Gems IV (Academic Press Professional, Inc.), pp. 474–485.

26. Klip, A., McGraw, T.E., and James, D.E. (2019). Thirty sweet years of GLUT4. J. Biol. Chem. 294, 11369–11381. 10.1074/jbc.REV119.008351.

27. Morris, S., Geoghegan, N.D., Sadler, J.B.A., Koester, A.M., Black, H.L., Laub, M., Miller, L., Heffernan, L., Simpson, J.C., Mastick, C.C., et al. (2020). Characterisation of GLUT4 trafficking in HeLa cells: comparable kinetics and orthologous trafficking mechanisms to 3T3-L1 adipocytes. PeerJ 8, e8751. 10.7717/peerj.8751.

28. Kylies, D., Zimmermann, M., Haas, F., Schwerk, M., Kuehl, M., Brehler, M., Czogalla, J., Hernandez, L.C., Konczalla, L., Okabayashi, Y., et al. (2023). Expansion-enhanced super-resolution radial fluctuations enable nanoscale molecular profiling of pathology specimens. Nat. Nanotechnol. 10.1038/s41565-023-01328-z.

29. Engdahl, A.K., Belle, S., Wang, T.-C., Hellmann, R., Huser, T., and Schuttpelz, M. Large Field-of-View Super-Resolution Optical Microscopy Based on Planar Polymer Waveguides.

30. Schindelin, J., Arganda-Carreras, I., Frise, E., Kaynig, V., Longair, M., Pietzsch, T., Preibisch, S., Rueden, C., Saalfeld, S., Schmid, B., et al. (2012). Fiji: an open-source platform for biological-image analysis. Nat. Methods 9, 676–682. 10.1038/nmeth.2019.

31. Preibisch, S., Saalfeld, S., and Tomancak, P. (2009). Globally optimal stitching of tiled 3D microscopic image acquisitions. Bioinformatics 25, 1463–1465. 10.1093/bioinformatics/btp184.

32. Roccisana, J., Sadler, J.B.A., Bryant, N.J., and Gould, G.W. (2013). Sorting of GLUT4 into its insulin-sensitive store requires the Sec1/Munc18 protein mVps45. Mol. Biol. Cell 24, 2389–2397. 10.1091/mbc.e13-01-0011.

33. Shannon, C.E. (1949). Communication in the Presence of Noise. Proc. IRE 37, 10– 21. 10.1109/JRPROC.1949.232969.

