## Supplementary Information for "Obtaining super-resolved images at the mesoscale through Super-Resolution Radial Fluctuations"

### Supplemental Information

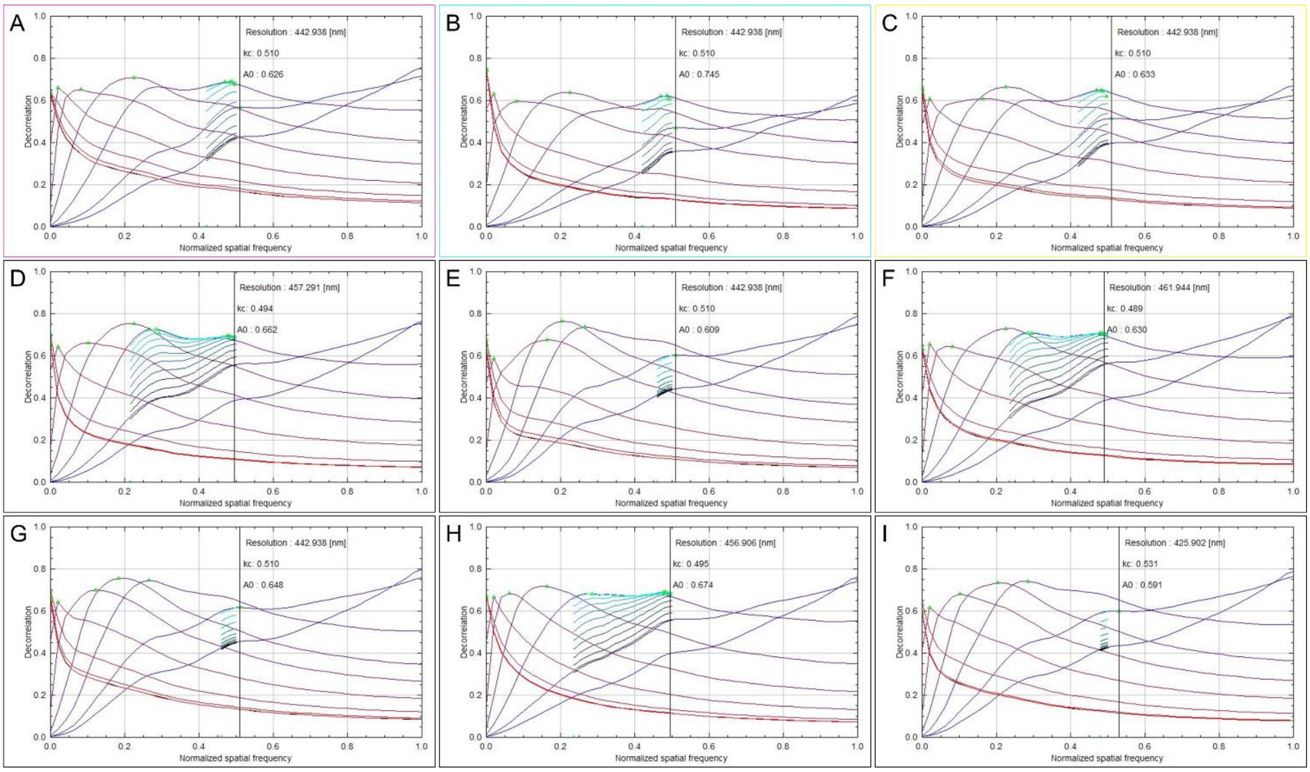


Figure S1: The plots of all decorrelation curves calculated from the ROIs highlighted in Figures 1 and 2, and the additional biological replicates used to calculate the achievable resolution.

(A-C) The decorrelation curves calculated from biological replicate 1, with the magenta, cyan, and yellow ROI as highlighted in Figures 1 and 2.

(D-F) The decorrelation curves calculated from three ROI from biological replicate 2.

(G-I) The decorrelation curves calculated from three ROI from biological replicate 3.

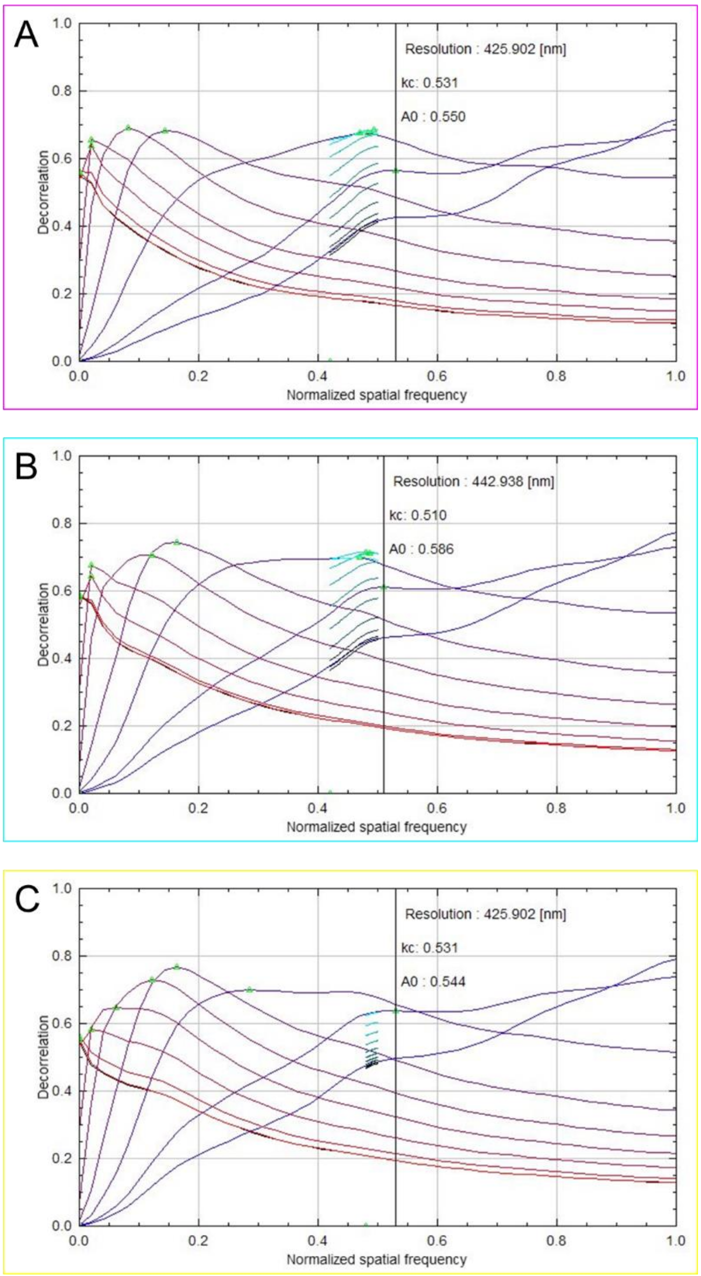


Figure S2: The plots of all decorrelation curves calculated from the ROIs highlighted in Figures 3 and 4 and used to calculate the resolution of GFP-tagged GLUT4 glucose transporters in 3T3-L1 fibroblasts.

(A) The decorrelation curves calculated from the magenta ROI.

(B) The decorrelation curves calculated from the cyan ROI.

(C) The decorrelation curves calculated from the yellow ROI.
